# Na-Dene populations descend from the Paleo-Eskimo migration into America

**DOI:** 10.1101/074476

**Authors:** Pavel Flegontov, N. Ezgi Altınışık, Piya Changmai, Edward J. Vajda, Johannes Krause, Stephan Schiffels

## Abstract

Prehistory of Native Americans of the Na-Dene language family remains controversial. Genetic continuity of Paleo-Eskimos (Saqqaq and Dorset cultures) and Na-Dene was proposed under the three-wave model of America’s settlement; however, recent studies have produced conflicting results. Here, we performed reconstruction and dating of Na-Dene population history, using genome sequencing data and a coalescent method relying on rare alleles (Rarecoal). We also applied model-free approaches for analysis of rare allele and autosomal haplotype sharing. All methods detected Central and West Siberian ancestry exclusively in a fraction of modern day Na-Dene individuals, but not in other Native Americans. Our results are consistent with gene flow from Paleo-Eskimos into the First American ancestors of Na-Dene, and a later less extensive bidirectional admixture between Na-Dene and Neo-Eskimos. The dated gene flow from Siberia to Na-Dene is in agreement with the Dene-Yeniseian language macrofamily proposal and with the succession of archaeological cultures in Siberia.

## Introduction

The Na-Dene language family includes the Tlingit, Eyak (recently extinct), Northern and Southern Athabaskan branches, and occupies Alaska and parts of Canada, with isolated groups residing along the US North Pacific Coast and much further south in the USA^1^. Under the three-wave model of America’s settlement, originally advanced in 1986 based on a synthesis of linguistic, dental and limited genetic data available at that time^2^, Na-Dene-speaking ethnic groups, further referred to as Na-Dene, were considered descendants of the second migration wave from Beringia. Genetic and archaeological data accumulated since 1986 have generally supported the three-wave model (reviewed by Skoglund and Reich^3^). The first major migration started about 16,000 years before present (YBP) and rapidly spread across North and South America^4^. We refer to the descendants of this migration as First Americans. The second, Paleo-Eskimo, migration associated with the Saqqaq and Dorset archaeological cultures, which are part of the Arctic Small Tool tradition, ASTt, took place about 4,800 YBP, long after the Bering land bridge had been inundated^5^. The third wave associated with the Thule culture occurred at about 1,000 YBP and gave rise to modern Yupik in Chukotka and Alaska and to Inuit throughout the American Arctic^6^. These ethnic groups are often referred to as “Eskimo”, and following Raghavan *et al.*^6^, we call this migration Neo-Eskimo. Both the second and the third waves were largely restricted to the American Artic. It was shown that genetic continuity characterized the Paleo-Eskimo period from about 4,800 to 700 YBP, and that Neo-Eskimos also maintained genetic continuity from the Siberian Old Bering Sea culture, 2,200 YBP, to modern Yupik and Inuit^6^.

While it is clear that the first and the third waves have left modern descendants, it remains controversial whether the second migration wave was completely replaced by the third one and whether it has left genetic traces in modern Na-Dene^3, 6-8^. Archaeologically, it is established that the latest Paleo-Eskimo cultures were replaced by the Thule culture, and it remains uncertain whether their contact and material exchange was extensive^6, 9, 10^. According to a recent reassessment of radiocarbon dating, the temporal overlap of Paleo- and Neo-Eskimos lasted 50 to maximum 200 years in the eastern American Arctic^6^.

Using single nucleotide polymorphism (SNP) array data for a large panel of Native American populations, Reich *et al.*^8^ have shown that Chipewyans, a Na-Dene-speaking Northern Athabaskan ethnic group, derive 16% of their ancestry from a population related to the 4,000-year-old Saqqaq ancient human genome from West Greenland^11^, and 84% of their ancestry is derived from First Americans related to Algonquins. A hypothesis that Northern Athabaskans derive their ancestry from admixture of First Americans and Neo-Eskimo populations has been rejected by a statistical test^8^. Other studies have challenged this conclusion^6, 7^. Based on genome-wide data from ancient Paleo- and Neo-Eskimos, modern Na-Dene and other Native Americans, the authors argued that: 1/ Dakelh, a Northern Athabaskan group from British Columbia, has no Paleo-Eskimo ancestry, but considerable admixture from Neo-Eskimos was detected; 2/ admixture of Saqqaq and Neo-Eskimo ancestors happened in Beringia prior to their entry into America; 3/ Neo-Eskimos completely replaced the Paleo-Eskimo population by about 1300 CE^6, 7^.

In order to shed light on these conflicting results, in this study we performed a detailed reconstruction and dating of the Na-Dene population history, using an extensive set of public sequencing data (Fig. 1) and a recently developed coalescent method, Rarecoal^12^, which relies on rare alleles and for which we developed a new extension to include admixture. In parallel, we generated new genotyping data from Siberian populations, and inferred and dated admixture events using GLOBETROTTER, a tool relying on autosomal haplotype data^13^. We also applied model-free approaches for analysis of rare allele and autosomal haplotype sharing.

**Fig. 1.**
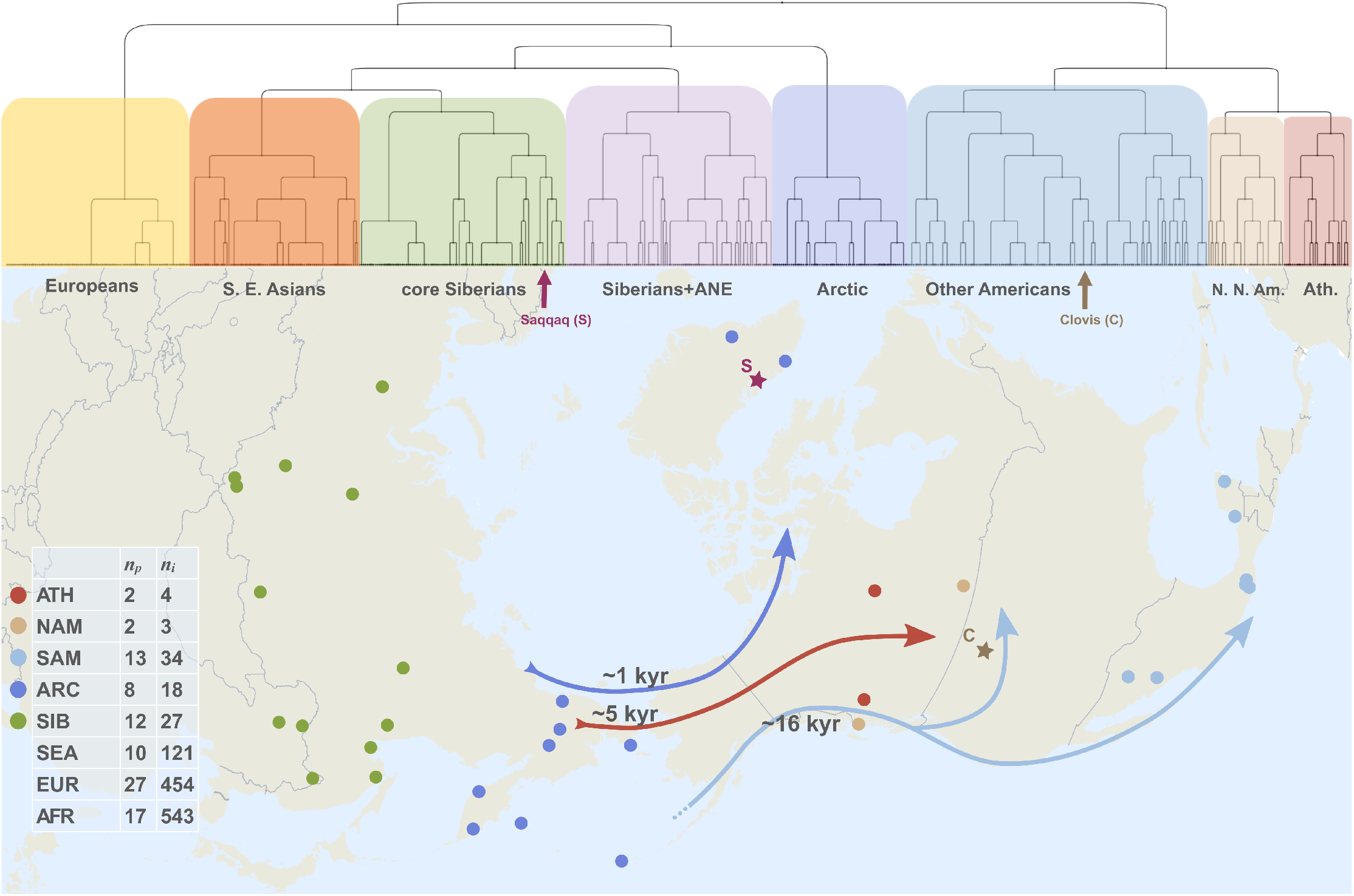
Genome sequencing data used for rare allele analysis. All data were taken from published sources, see Suppl. Table 1. The dataset composition (number of populations, *n*_*p*_, and individuals, *n*_*i*_, in each meta-population) is shown in the table on the left. Meta-populations are color-coded in a similar way throughout all figures and designated as follows: Arctic (abbreviated as ARC); Na-Dene, in this analysis represented by two Northern Athabaskan groups, Chipewyan and Dakelh (ATH or Ath.); Europe and the Caucasus (EUR); northern North Americans, excluding Na-Dene, Yupik and Inuit (NAM or N. N. Am.); Southeast Asians (SEA or S. E. Asians); Siberians, excluding populations of Chukotka and Kamchatka (SIB); native populations of South, Central America, Mexico and southern USA (SAM). Locations of Siberian, Arctic, Athabaskan, and North American populations are shown on the map below, which also illustrates three migration waves and their approximate dates in thousands of year (kyr). Locations of the Saqqaq (dated at about 4,000 YBP) and Clovis (12,600 YBP) ancient genomes are shown with asterisks. On top a clustering tree constructed with fineSTRUCTURE on aversion of HumanOrigins SNP array dataset (655 individuals and 58 populations, see Suppl. Table 2) illustrates relationships of the meta-populations. For a detailed version of the same tree see Suppl. Fig. 3A.

## Results

### Dataset composition and sample information

We composed a large set of sequencing data covering Africa, Europe, Southeast Asia, Siberia, and the Americas: 1,206 individuals from 94 populations, including the Clovis^14^ and Saqqaq^11^ ancient genomes (Fig. 1, Suppl. Table 1). For the purpose of haplotype sharing analysis, we composed two independent SNP array datasets covering the same geographic regions (Suppl. Table 1): 1/ a set based on the HumanOrigins platform (1,283 individuals from 101 populations, including Saqqaq and Clovis); 2/ a set based on various Illumina arrays (645 individuals from 63 populations, including Saqqaq). Properties of the datasets and their versions used for various analyses are described in Suppl. Table 2. We also present new data, from 58 Siberian individuals (Kets, Nganasans, Selkups, and Enets), which were genotyped for up to 612,164 autosomal SNPs on the HumanOrigins array platform (see sample information in Suppl. Table 3).

For rare allele sharing analyses relying on genome sequencing data, we combined ethnic groups into non-overlapping meta-populations (Suppl. Table 1), sharing a common genetic background: 1/ Sub-Saharan Africans; 2/ ethnic groups of Europe and the Caucasus, excluding populations mixed with Central Asians or Siberians (see Methods); 3/ Southeast Asians; 4/ the Arctic group - peoples of Beringia and the American Arctic that are derived from the third, Neo-Eskimo, wave of America’s settlement and from populations closely related to its Siberian source^6^; 5/ Siberians, excluding any populations of the previous group; 6/ Na-Dene ethnic groups; 7/ a group of northern North Americans, other than Na-Dene, which are genetically distinct from populations further to the South^3, 7, 8^; 8/ native populations of South, Central America, Mexico and southern USA^3, 7, 8^. For haplotype sharing analyses relying on genotyping data with larger population sizes, a more fine-grained breakdown was possible (Suppl. Table 1). The Arctic group was split into the Siberian and American Arctic subgroups; Siberians with extensive ancient North Eurasian ancestry^15, 16^ were considered separately and referred to as Siberians+ANE, while other Siberian groups were called ‘core Siberians’. The breakdown of ethnic groups into these meta-populations was supported by ADMIXTURE analysis on unlinked SNPs (Suppl. Fig. 1A,B) and by the principal component analysis (PCA, see Suppl. Fig. 2A,B) and clustering trees (Fig. 1, Suppl. Fig. 3A,B) constructed by fineSTRUCTURE^17^ based on autosomal haplotype sharing patterns. Meta-populations most relevant for our study are the following: Na-Dene with 4 high-coverage genomes, 32 and 48 individuals in the HumanOrigins and Illumina SNP array datasets, respectively; northern North Americans (3 genomes, 34 and 65 genotyped individuals); other Americans (34 genomes, 151 and 69 genotyped individuals); Arctic (18 genomes, 74 and 56 genotyped individuals) and Siberian groups (27 genomes, 221 and 190 genotyped individuals).

### Na-Dene stand out from the other Native Americans

To measure the relationship of Native American populations with Siberian and Arctic groups, we looked for rare SNP alleles shared between them. Rare variants, i.e. those occurring at a global frequency of less than 1%, have been shown to have more power to resolve subtle relationships than common variants^12,18^. We calculated allele sharing counts (ASC) and their standard deviations for each American population (a modern group or an ancient genome) and the Siberian or Arctic meta-populations (Fig. 2, see Methods for details). To take care of variability in genome coverage across populations and of dataset-specific SNP calling biases, we normalized counts of alleles shared between an American group and Siberian or Arctic meta-populations by similar counts of alleles shared with a distant outgroup. Europeans or Africans were used as alternative outgroups in this study. Since we expected a decay of a recent ancestry signal at higher allele frequencies^12^, all statistics were calculated separately for alleles of various frequencies: occurring 2, 3, 4, … and up to 20 times among 2,412 haploid chromosome sets, which corresponds to frequencies from 0.08% to 0.83%.

**Fig. 2.**
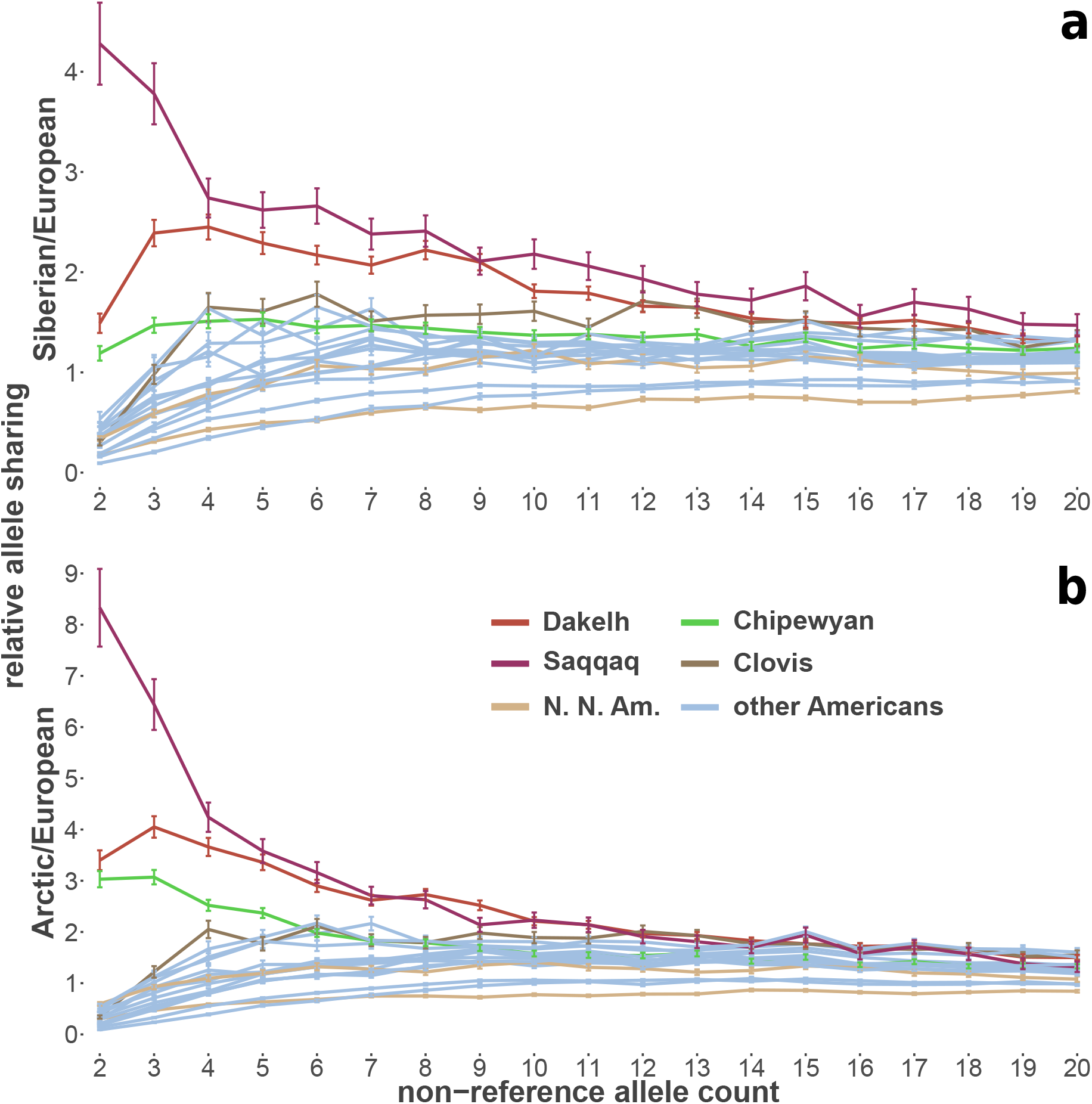
Relative rare allele sharing counts and their standard deviations calculated for each American population or ancient genome and the Siberian (**A**) or Arctic (**B**) meta-populations. All statistics were calculated separately for alleles of various frequency: occurring 2, 3, 4, … and up to 20 times in the set of 2,412 chromosomes. To take care of variability in genome coverage across populations and of dataset-specific SNP calling biases, we normalized the counts of alleles shared by a given American population and the Arctic or Siberian meta-populations by similar counts of alleles shared with distant outgroups -Europeans (this figure) or Africans (Suppl. Fig. 4A,B). Saqqaq and Northern Athabaskans (Chipewyans and Dakelh) stand out from northern North Americans (N. N. Am.) and other First American populations.

The Saqqaq ancient individual and Northern Athabaskans (Chipewyans and Dakelh) clearly stand out from other Americans, according to both Siberian and Arctic relative allele sharing (Fig. 2). This result is expected for Saqqaq, since its close relationship to the Arctic and Siberian groups was shown by various methods^6, 7, 11, 15^. We also note that in our analysis the 12,600-years-old Clovis ancient genome^14^ does not differ from modern South and Central American populations. The same results were obtained with Africans as the normalizer (Suppl. Fig. 4A-B), and when only private (i.e. exclusively shared between two meta-populations) alleles were counted (Suppl. Fig. 4C-F). As expected, privately shared alleles were largely restricted to the lowest-frequency bins: allele counts from 2 to 5, corresponding to frequencies from 0.08% to 0.21%.

We note that the Siberian and Arctic signals are stronger in the Dakelh genomes (two individuals^6,7^) as compared to the Chipewyan genomes (two individuals^19^), although both populations are significantly different from other Americans at low allele frequencies. For Dakelh, the signal is observed for allele counts up to 10 (0.42% frequency, see Fig. 2, Suppl. Fig. 4A,B), which suggests relatively recent gene flow^12^ from Siberian and/or Arctic populations into Northern Athabaskans. We investigate the source of this gene flow and attempt its dating in the following sections.

### Na-Dene belong to the Paleo-Eskimo wave of America’s settlement

While we have shown that Northern Athabascans have elevated Siberian and Arctic rare allele sharing compared to all other Native Americans investigated, this does not immediately suggest that they descend from the second, PaleoEskimo, settlement wave^8^ vs. the third, Neo-Eskimo, one^6,7^. Therefore, we combined the Siberian and Arctic allele sharing statistics on a two-dimensional plot showing each American population (Fig. 3). For that purpose, we summed up allele sharing statistics for allele counts from 2 to 5 – those demonstrating most prominent signals (Suppl. Fig. 4). For comparison, we also generated the same statistics with a take-one-out procedure for Siberians and for representatives of the American Arctic group (Fig. 3). We observe that each meta-population is scattered along a line on the plot, which reflects similar ratios of the Siberian and Arctic allele sharing among its members (Fig. 3A). The position of a population along the line depends on the presence of other ancestry components. For example, Aleuts, having a high level of European admixture^6, 7^ (see also Suppl. Fig. 1), lie much closer to zero as compared to the other American Arctic groups (Fig. 3B). While First American populations form a tight cluster, the Dakelh and Chipewyan populations are shifted considerably towards the Saqqaq individual. Since allele sharing counts behave linearly under admixture, we used linear combinations to calculate expected relative allele sharing statistics for recently admixed populations: mixtures of First Americans with either the Saqqaq individual or populations of the third migration wave. We used all modern First Americans, the Greenlander Inuit and two Chukotkan Yupik third-wave populations for this simulation, and assumed 70%, 75%, … and 90% of First American ancestry in the admixed populations.

**Fig. 3.**
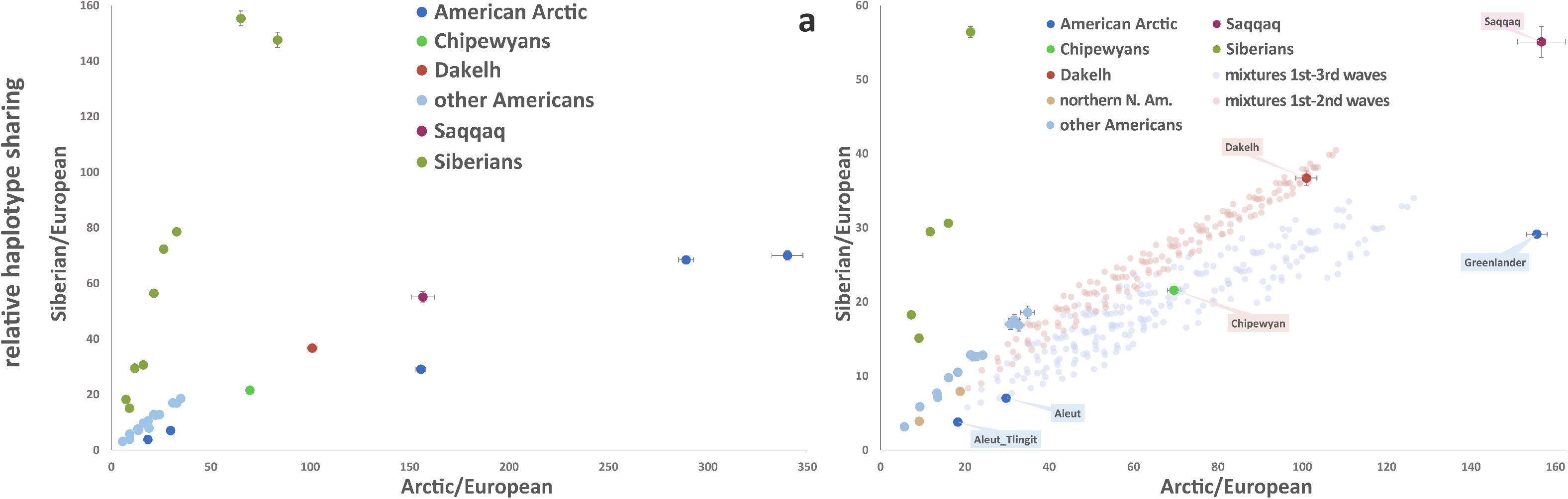
Two-dimensional plots of Siberian and Arctic allele sharing counts normalized using the European meta-population. (**A**) A plot showing statistics for all populations and standard deviations. Meta-populations are color-coded according to the legend. (**B**) An enlarged area of the plot showing simulated mixtures of any modern First American population and the Saqqaq ancient individual (from 10% to 60%), and similar mixtures with any third-wave population (from 10% to 30% of Greenlander Inuit or Chukotkan Yupik ancestry). Some population names are indicated on the plot.

Normalized allele sharing counts for the Dakelh population match those for a mixed Saqqaq/First American population and are clearly different from those of any simulated Neo-Eskimo/First American mixtures, i.e. are separated from the latter cluster by more than three standard error intervals (Fig. 3B). This result is consistent with Paleo-Eskimos being ancestors of Na-Dene. However, the Chipewyan population lies within the cluster of the first and third wave mixtures. Using Africans for the normalization (Suppl. Fig. 5A,B,E,F) and/or counting private alleles gives similar results (Suppl. Fig. 5C-F).

To investigate a wider diversity of Na-Dene and other northern North American populations, we applied a similar analysis strategy to a different type of data: to autosomal haplotype sharing statistics on the HumanOrigins and Illumina SNP array datasets (Suppl. Tables 1 and 2). Cumulative lengths and counts of shared autosomal haplotypes were produced with ChromoPainter v.1 for pairs of individuals, in the form of all vs. all “coancestry matrices”^17^, then American-Siberian or American-Arctic haplotype sharing statistics were calculated for each American individual and normalized using a distant outgroup (see Methods for details). Two-dimensional plots showing Saqqaq, Na-Dene, and other relevant meta-populations appear in Fig. 4 and Suppl. Fig. 6 for the HumanOrigins dataset, and for the Illumina dataset in Suppl. Figs. 7 and 8.

**Fig. 4.**
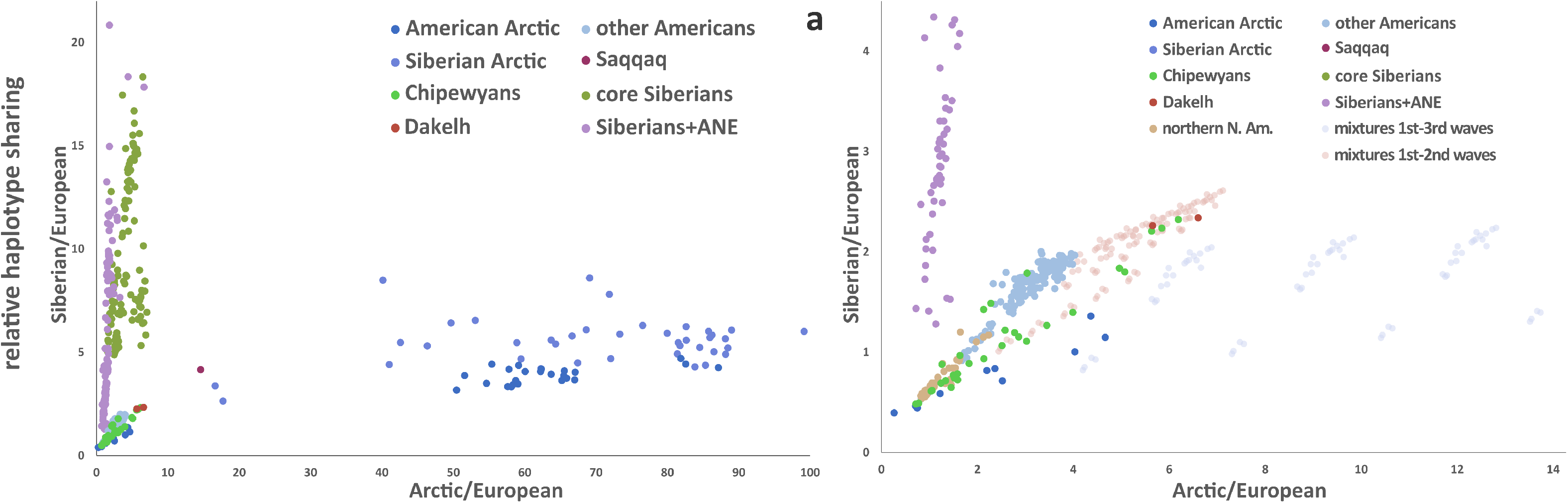
Two-dimensional plots of Siberian and Arctic haplotype sharing statistics normalized using the European meta-population. (**A**) A plot showing statistics for individuals of all relevant meta-populations, color-coded according to the legend. (**B**) An enlarged area of the plot showing statistics for American individuals and simulated mixtures of any modern First American population and the Saqqaq ancient individual (from 10% to 30%), and similar mixtures with the Chukotkan Yupik (Eskimo) population (from 5% to 20% of Yupik ancestry). Average values of the statistics in populations were used to calculate the simulated statistics.

Most northern North American ethnic groups have post-Columbian European admixture, highly variable among individuals^7,20^. The same pattern was observed in both of our datasets with ADMIXTURE (Suppl. Fig. 1), and in the Illumina dataset a number of northern North American and Na-Dene individuals formed a clade with Europeans, while others clustered with South Americans (Suppl. Fig. 3B). Moreover, the Siberian and Arctic groups have considerable European admixture, dated to 2,200-2,400 YBP in the former (see Table 1 and Suppl. Text 1). Thus, we expected the post-Columbian European ancestry in northern North Americans and Na-Dene to bias the Siberian and Arctic haplotype sharing statistics upwards. To mitigate this potential bias, we preferred to use haplotype sharing with Europeans as a normalizer and plotted statistics for individuals, since we expected highly variable levels of European ancestry in North Americans. Essentially similar results were produced on both SNP array datasets, using both European and African normalizers (Fig. 4, Suppl. Figs. 6, 7, 8).

**Table 1.** Final parameter estimates including posterior probability distribution quantiles for the six-population demographic model with asymmetric migration. Meta-populations are abbreviated as follows: American Arctic (AARC), Athabaskan (ATH), European (EUR), northern North American (NAM), South American (SAM), Southeast Asian (SEA), Siberian (SIB). See Suppl. Text 1 for further details on populations used in the analysis. The right-most column shows parameters for a model where the Athabaskan group was replaced by northern North Americans. Parameters most important for our discussion are shaded in grey.

|  | Parameter | Maximum Likelihood estimate (ATH) | 2.5% posterior quantile | 50% posterior quantile (Median) | 97.5% posterior quantile | Maximum Likelihood estimate (NAM) |
| --- | --- | --- | --- | --- | --- | --- |
| **effective population sizes** | EUR | 25,101 | 25,015 | 25,094 | 25,182 | 25,714 |
|  | SEA | 44,242 | 43,943 | 44,347 | 44,720 | 44,620 |
|  | SIB | 13,568 | 13,115 | 13,445 | 13,710 | 10,303 |
|  | AARC | 1,173 | 1,149 | 1,177 | 1,193 | 780 |
|  | ATH | 1,851 | 1,803 | 1,847 | 1,890 | 5,280 |
|  | SAM | 6,552 | 6,485 | 6,589 | 6,717 | 5,664 |
|  | EUR-SEA... | 9,315 | 9,272 | 9,316 | 9,359 | 9,341 |
|  | SEA-SIB... | 9,012 | 8,961 | 9,031 | 9,106 | 8,753 |
|  | SIB...-SAM... | 147 | 146 | 148 | 149 | 141 |
|  | SAM-ATH | 1,762 | 1,750 | 1,765 | 1,783 | 1,610 |
|  | SIB-AARC | 27,469 | 27,202 | 27,621 | 28,003 | 28,612 |
| **split times** | EUR-SEA... | 36,095y | 35,980y | 36,131y | 36,279y | 35,588y |
|  | SEA-SIB... | 20,402y | 20,373y | 20,410y | 20,450y | 20,374y |
|  | SIB...-SAM... | 20,290y | 20,217y | 20,253y | 20,289y | 20,271y |
|  | SAM-ATH | 9,744y | 9,591y | 9,714y | 9,829y | 11,792y |
|  | SIB-AARC | 4,126y | 4,011y | 4,116y | 4,195y | 2,580y |
| **admixture proportions and dates** | EUR → SIB | 16.1% | 15.9% | 16.1% | 16.3% | 16.6% |
|  | SIB → EUR | 8.0% | 7.9% | 8.0% | 8.2% | 6.7% |
|  | date EUR ↔ SIB* | 2,327y | 2,198y | 2,299y | 2,402y | 1,753y |
|  | EUR → AARC** | 25.0% | 24.8% | 25.0% | 25.2% | 18.8% |
|  | EUR → SAM** | 2.7% | 2.6% | 2.7% | 2.7% | 3.1% |
|  | EUR → ATH** | 0.7% | 0.4% | 0.7% | 1.0% | 29.2% |
|  | SIB → ATH | 22.9% | 22.3% | 23.0% | 23.8% | 9.6% |
|  | ATH → SIB | 6.8% | 6.5% | 6.8% | 7.0% | 1.6% |
|  | date SIB ↔ ATH* | 6,940y | 6,575y | 6,812y | 7,030y | 595y |
|  | AARC → ATH | 7.6% | 6.3% | 7.5% | 8.5% | 4.8% |
|  | ATH → AARC | 11.5% | 10.9% | 11.6% | 12.4% | 17.4% |
|  | date AARC ↔ ATH* | 490y | 476y | 492y | 499y | 5y |
|  | SAM → AARC | 7.6% | 7.2% | 7.4% | 7.7% | 21.9% |
|  | date SAM → AARC | 488y | 473y | 481y | 495y | 2,570y |

\* To reduce the overall number of parameters, admixture events in both directions were enforced to occur at the same time.
\*\* For the same purpose, European admixture in Americans (AARC, ATH, NAM, and SAM groups) was modeled as unidirectional with the age of 500 years.

Haplotype sharing statistics in both Dakelh individuals and in a fraction of Chipewyans (3 of 30) match those of simulated First American populations having from ~20% to ~30% of Saqqaq ancestry, and are different from those of any First American/Neo-Eskimo mixtures. Ten Chipewyan individuals are less shifted from the cluster of First American individuals, and their haplotype sharing statistics might be explained either by a proportion of Saqqaq ancestry of about 10-20% or by very low levels of Chukotkan Yupik ancestry (<<5%, see Fig. 4B). Besides Northern Athabaskans, other major branches of the Na-Dene language family were represented in the Illumina SNP array dataset, namely Southern Athabaskans from USA and Tlingit from western Canada (Suppl. Table 1). Four Dakelh, three other Northern Athabaskans, six Tlingit, one Southern Athabaskan, and only one non-Na-Dene individual (1 of 7 Splatsin) showed a signal of Paleo-Eskimo (Saqqaq) admixture (Suppl. Fig. 7B). Notably, one Northern Athabaskan individual (population ‘Northern Athabaskan 3’ according to Raghavan *et al.*^7^) corresponded to a ~30% mixture of Saqqaq and ~70% of First Americans (Suppl. Fig. 7B). A third of Northern Athabaskans (7 of 20) showed a pronounced signal of Paleo-Eskimo admixture, while most investigated Tlingit (17 of 23) and Southern Athabaskans (4 of 5) did not show a clear signal. However, haplotype sharing statistics of few other Na-Dene and northern North American individuals are compatible with low levels of Paleo-Eskimo (~10%) or Inuit ancestry (~5%) (Suppl. Fig. 7B). Of the four Dakelh with a signal of Paleo-Eskimo ancestry, two individuals had corresponding genome sequencing data^6^, and they have demonstrated consistent results throughout all analyses: on sequencing data (Fig. 3, Suppl. Fig. 5), on sequencing data merged with the HumanOrigins SNP array (Fig. 4, Suppl. Figs. 3A, 6), and on Illumina SNP arrays (Suppl. Figs. 3B, 7, 8).

In summary, our model-free approach to analyze rare allele and haplotype sharing reveals that a fraction of Na-Dene Native Americans likely has a considerable proportion of Paleo-Eskimo ancestry, roughly from 10 to 30%. Virtually no other Native Americans demonstrated the same signal in our analysis, despite a large number of populations and individuals investigated (37 genomes and 319 genotyped First Americans).

### Demographic modeling and dating of population mixtures

To interpret our findings in a more quantitative way, we built an explicit demographic model for the peopling of North America. We used Rarecoal^12^ to estimate split times and population sizes, as well as admixture events, in a population tree connecting Europeans, Southeast Asians, Siberians, populations of the Neo-Eskimo migration wave, Northern Athabaskans, and Native South Americans. Sample sizes and additional details are provided in Suppl. Text 1. Rarecoal is a software that implements a fast algorithm to estimate the joint site frequency spectrum for rare alleles in hundreds of samples^12^. Since the initial report, we have improved the software and added pulse-like admixture events as a new feature (see Methods).

The model was derived in an iterative way: we started off with fitting a model to three populations only (Europeans, Southeast Asians, and South Americans), and then added one population at a time, re-estimating all previous and new parameters (see details in Suppl. Text 1). Admixture edges were added when the model fit showed significant deviations for particular allele sharing statistics. The final model (Fig. 5A, Table 1) contains six clades, four unidirectional admixture edges and three bidirectional edges with asymmetric admixture rates. The parameter estimates including confidence intervals for this final model are shown in Table 1.

**Fig. 5.**
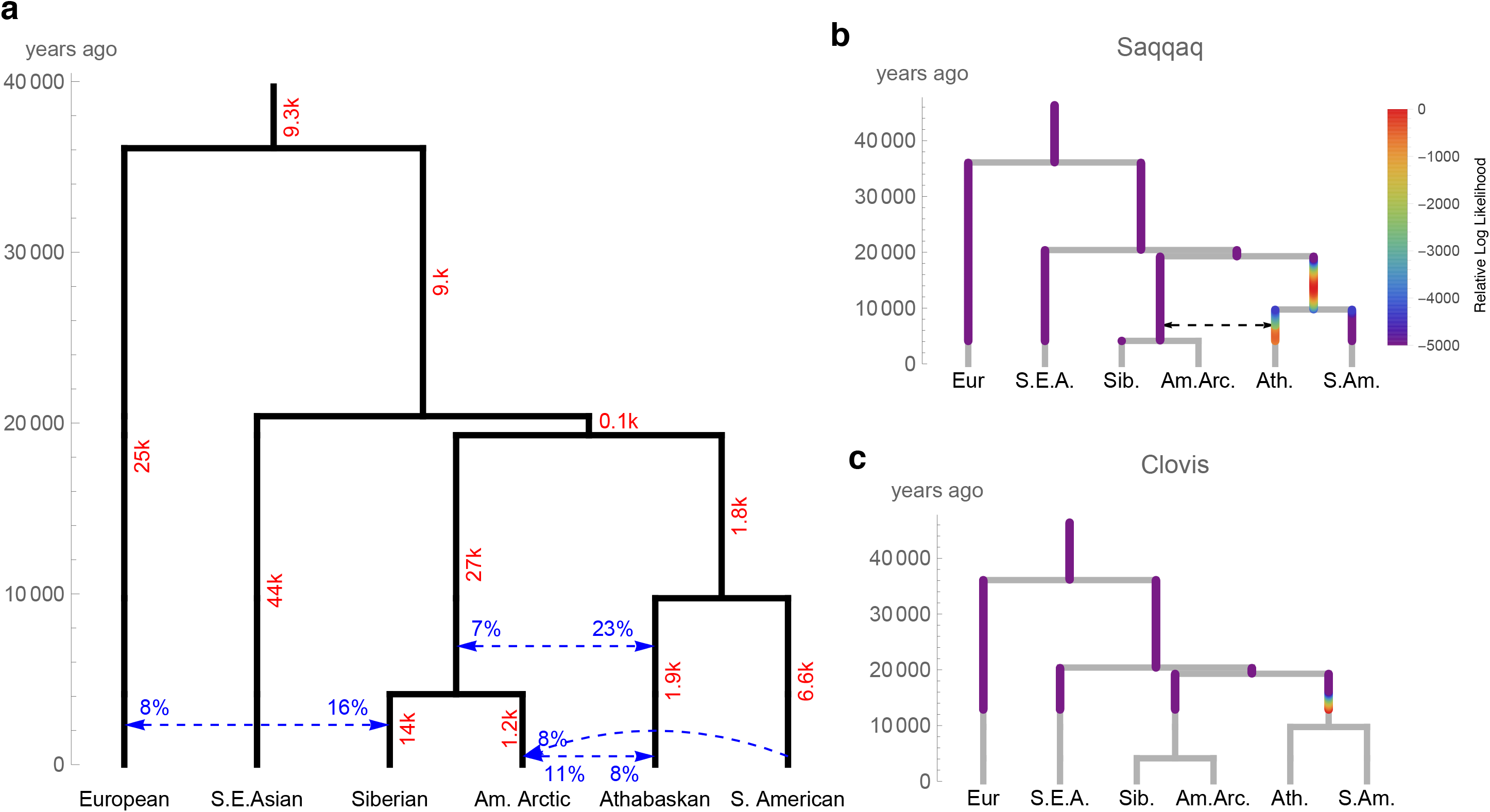
A dated six-population demographic model with asymmetric migration constructed using Rarecoal. For a complete list of parameter estimations see Table 2. Meta-populations are abbreviated as follows: American Arctic (Am.Arc.), Athabaskan (Ath.), European (Eur.), South American (S.Am.), Southeast Asian (S.E.A.), Siberian (Sib.). (**A**) For three edges most important for our study (European-Siberian, Siberian-Athabaskan, Athabaskan-American Arctic), separate estimations of gene flow in both directions were performed. To reduce the overall number of parameters, these admixture events were enforced to occur at the same time. For the same purpose, European admixture in Americans (in the American Arctic, Athabaskan and South American groups) was modeled as unidirectional with the age of 500 years, and these edges are omitted for clarity (see their parameters in Table 2). Effective population sizes (in 1000) are shown in red. Most likely branching points for the Saqqaq (**B**) and Clovis (**C**) ancient genomes were also estimated using Rarecoal.

Substantial admixture of 22.3 – 23.8% from Siberians (22 genomes) into Northern Athabaskans was revealed in our model, with only 6.5 – 7% in the opposite direction (95% confidence intervals are given). The Siberian-Athabaskan admixture edge was dated at 6,575 – 7,030 YBP (Table 1). A simpler model without the American Arctic meta-population (Suppl. Text 1) dated the Siberian-Athabaskan admixture at ~4,400 YBP, which likely corresponds to the admixture of Paleo-Eskimos and First Americans that must postdate the Paleo-Eskimo immigration around 4,800 YBP^5^. Admixture was also inferred between the American Arctic groups and Athabaskans, however, with a much smaller admixture proportion of 6.3 – 8.5% into Athabaskans and 10.9 – 12.4% in the opposite direction, and a much later date of 476 – 499 YBP (Table 1). We caution that the assumption of constant population sizes within branches, which is necessary to keep the number of parameters manageable, may lead to overly narrow confidence intervals of our estimates.

In order to assess whether the Siberian admixture inferred in Athabaskans is also present in other northern North Americans, we tested the final model shown above on a data set where Athabaskans are replaced with non-Na-Dene speaking Cree (2 genomes) and Tsimshian (1 genome). On this data, we still estimate ~10% Siberian admixture into northern North Americans (compare with 23% from Siberians into Athabaskans). However, the time of this admixture event (~600 YBP) is extremely recent, and moreover after the European admixture event into Siberians (Fig. 5A, Table 1). We think that this may reflect recent admixture between Athabaskans and other Northern Americans. In any case, the signal is weaker and too recent to reflect the same historical admixture event that is seen in the Athabaskans.

To test the robustness of our estimates we simulated the final six-population model with the Athabaskans under the coalescent with recombination (see Suppl. Text 1). We then estimated parameters from the simulated data using Rarecoal and checked whether the inferred parameters match the simulated parameters. The results are summarized in Suppl. Fig. 1.15. As can be seen, most parameters are estimated very accurately, in particular all time estimates of splits and admixture events. Substantial deviation between a simulated and estimated parameter is seen in the population size estimate of the Siberian branch, as well as the ancestral branch of Siberians and the American Arctic groups. This could reflect substructure in our Siberian and/or American Arctic meta-populations, which was not part of the simulation but could have an effect on model estimates.

Finally, we attempted to map the high-coverage Saqqaq and Clovis ancient genomes onto the modeled tree. It is hardly possible to incorporate single individuals fully into the model, and low sequencing coverage of other Paleo-Eskimo genomes available^6^ makes them much less suitable for our analysis. Instead, we evaluated the likelihood of a sample’s branch to merge onto the tree, testing all time points on all branches, before the age of the sample. The Saqqaq genome^11^ most likely branches off the tree either before the split of Athabaskans and South Americans, or at the Athabaskan branch immediately after the gene flow from Siberians (Fig. 5B). The branching point of the 12,600-years-old Clovis genome^14^ fits its expected position at the base of the American clade (Fig. 5C).

The somewhat surprising clustering of the Saqqaq genome onto the Native American ancestral branch in our Rarecoal analysis may reflect subtle differences between Saqqaq and the extant Siberians and American Arctic populations used for constructing the model (Suppl. Text 1). In all previous analyses^6, 11^, ^15^ and in haplotype-based analyses in this study (Fig. 1, Suppl. Fig. 3B), Saqqaq clustered with either core Siberian or Siberian Arctic groups, probably reflecting the fact that it branched off the Siberian stem prior to the separation of the modern groups (the Chukchi-Saqqaq divergence was dated at 4,400 — 6,400 YBP^11^). The branch point inferred by Rarecoal probably reflects a Siberian ancestor of Saqqaq, that is closer to Native American ancestors than to the ancestors of the Siberian and American Arctic samples used here. The other high-likelihood branch point for Saqqaq, on the Athabaskan branch after the Siberian admixture event, suggests that the Siberian-Athabaskan gene flow modeled here was mediated by Paleo-Eskimos. In any case, Paleo-Eskimos represent the most likely vector for any relatively recent gene flow from Siberia that pre-dates the Neo-Eskimo migration around 1,000 YBP, since no other ancient American group has been shown to possess detectable levels of “core Siberian” ancestry^6^.

Substantiating this conclusion, an admixture event between Saqqaq and First Americans was revealed in the history of Northern Athabaskans using GLOBETROTTER, a haplotype-based tool capable of inferring and dating up to two distinct admixture events^13^. GLOBETROTTER operates on coancestry curves, generated from ChromoPainter v.2^13^ (see Methods for details). In our analysis, two-date curves fit the Na-Dene data better as compared to one-date curves (Table 2, Suppl. Fig. 9). GLOBETROTTER also finds the closest proxies of admixture partners in a given dataset and determines admixture ratios. Saqqaq and First Americans were revealed as most likely admixture partners, with Saqqaq contribution in the 19% — 25% range, depending on a dataset (see Table 2), in good agreement with the Rarecoal results above. Using meta-populations as haplotype donors and five Northern Athabaskans we expected to be admixed with Paleo-Eskimos (Fig. 4), the Saqqaq admixture was dated at ~3,600 YBP with a 95% confidence interval of 488 — 4,614 YBP (Table 2).

**Table 2.**
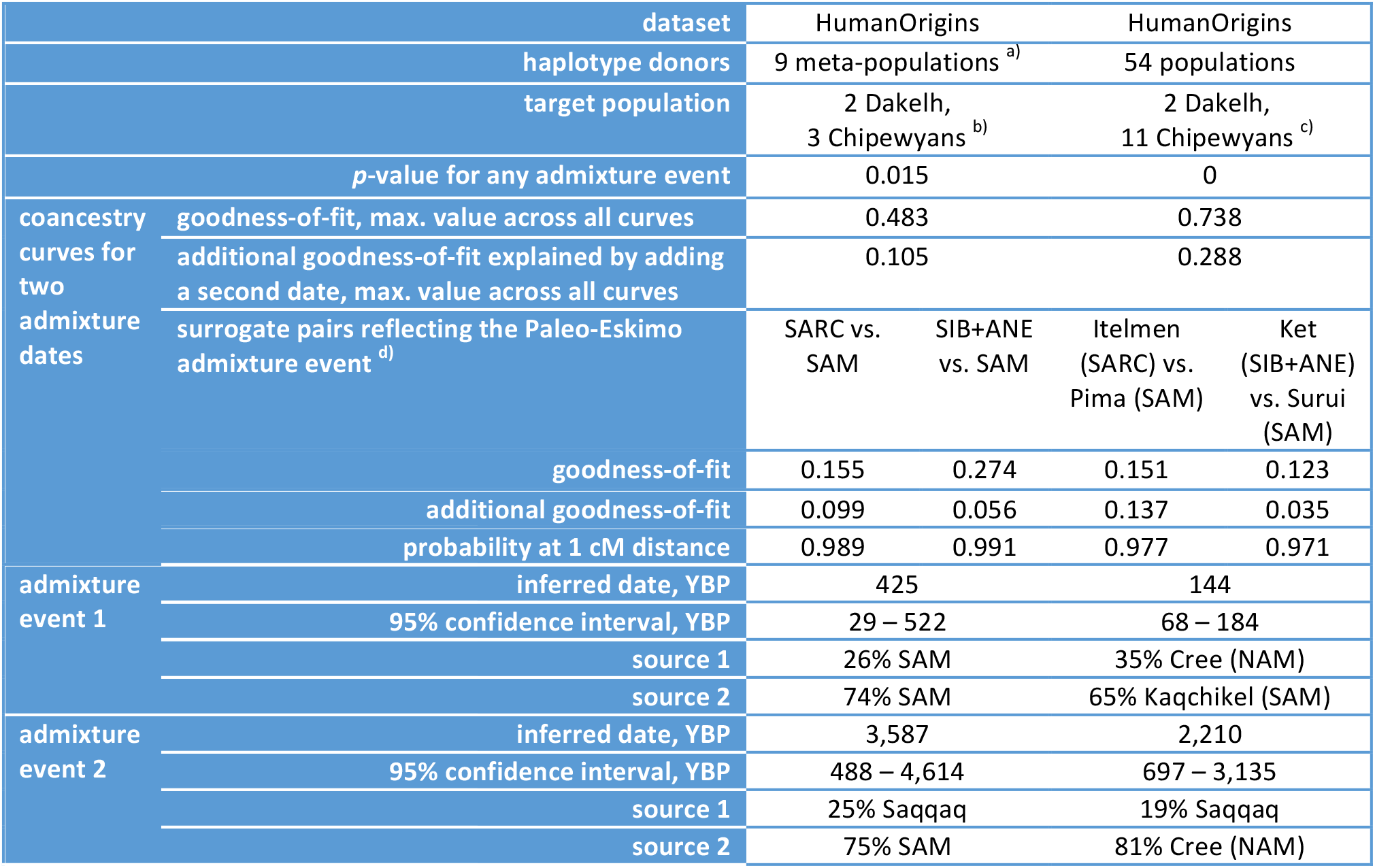
Dating admixture events in the history of Na-Dene populations using haplotype sharing data, generated with ChromoPainter v.2 and analyzed with the GLOBETROTTER approach^13^. Coancestry curves approximating two distinct admixture dates have always demonstrated a better fit to the data (Suppl. Fig. 9), and fit statistics for these curves are shown in this table, as well as inferred mixture partners, mixture proportions, dates and their 95% confidence intervals.

## Discussion

Our results are consistent with a gene flow from the Saqqaq Paleo-Eskimos (19 -25% admixture ratio) exclusively into the First American ancestors of Na-Dene, and a much later and less extensive bidirectional gene flow was detected between the Na-Dene and Neo-Eskimo branches. A somewhat lower level of Saqqaq-related ancestry of 16% was reported using the admixture graph method in Chipewyans, a Northern Athabaskan ethnic group^8^. We emphasize that only a fraction of modern Na-Dene individuals displays this level of Saqqaq ancestry, with most Na-Dene being admixed with other native groups and/or Europeans.

Methods of rare allele and autosomal haplotype analysis are especially sensitive for reconstructing recent population history within a few thousand years, and in some cases were demonstrated to outperform traditional methods^12, 13, 30^ based on unlinked common genetic variants, such as ADMIXTURE, PCA, TreeMix, *f*_*3*_-, *f*_*4*_-, and *D*-statistics. In this light, we consider discrepancies between our results and those of previous studies in Suppl. Text 2.

We dated the Paleo-Eskimo admixture at about 3,600 YBP using GLOBETROTTER analysis based on autosomal haplotypes (Table 2). Much older dates of the Siberian-Athabaskan gene flow obtained in our analysis based on rare allele sharing, about 6,500-7,000 YBP (Fig. 5, Table 1), probably correspond not to the admixture of Paleo-Eskimos and First Americans (that must postdate the Paleo-Eskimo immigration), but to a time point when Paleo-Eskimo ancestors branched off from the Siberian-Arctic stem. The split date suggested here for this unsampled “ghost population” fits the archaeological record of Siberia remarkably well, as discussed below. And split dates for other nodes inferred here broadly agree with the dates produced with independent methods^4, 7^.

The new wave of population from northeastern Asia that arrived in Alaska at least 4,800 years ago^5^ displays clear archaeological precedents leading back to Central Siberia. The rise of the Syalakh culture that flourished across much of Northeastern Siberia between 6,500 and 5,200 YBP involved migrants from the Transbaikal area who possibly mixed with local remnants of the earlier Sumnagin culture (10,500-6,500 BP), bringing the bow and arrow and new types of pottery to Northeastern Siberia^21, 22^. As the Bel’kachi culture (5,200-4,100 YBP) developed from Syalakh along the Lena and Aldan rivers^23^, at least one group of these people might have crossed the Bering Strait into Alaska around 4,800 YBP^5^, giving rise to Paleo-Eskimos. Thus, the Syalakh culture peoples, spreading across Siberia after 6,500 YBP, might represent the “ghost population” that split off around 6,500-7,000 YBP and later gave rise to migrants into America.

The geographic connection between Paleo-Eskimos and the related Siberian groups probably became severed as subsequent waves of hunter-gatherers entering Eastern Siberia from the west during the Late Neolithic (Ymyakhtakh culture, 3,700-2,800 YBP) brought new cultures and new language groups^24^. This phase of North Asian prehistory most likely involved the spread of Yukaghir, Chukchi-Kamchatkan and Eskimo-Aleut languages^25^, whose presence in the extreme northeast of Asia intervened geographically between Paleo-Eskimos, Na-Dene, and their Old World cousins. Notably, the dates of the Siberian-Arctic split obtained under our model (~4,000-4,200 YBP, Table 2) also agree with this scenario that links the spread of the Ymyakhtakh culture (after 3,700 YBP) with the Arctic meta-population, i.e. ancestors of modern Chukchi-Kamchatkan and Eskimo-Aleut ethnic groups.

The success of Paleo-Eskimos and Na-Dene in occupying territories previously populated by First Americans^5, 28^, in some cases (Southern Athabaskans) moving very far from the original homeland in Alaska and northwestern Canada, might be partially attributed to archery, a technological advance lacking among the local populations. Paleo-Eskimos quickly spread from Alaska to Greenland and Labrador and have been credited with introducing the bow and arrow to populations in Eastern Canada by 4,000 YPB^29^, though the Dorset people, the last wave of Paleo-Eskimos, seem to have given up this technology for handheld lances^10^.

Another important observation concerns the distribution of Siberian (Paleo-Eskimo) ancestry among modern North Americans. The methods used in this study detected Central and West Siberian ancestry in a fraction of Na-Dene individuals belonging to all major branches of the language family existing today: Tlingit, Northern Athabaskan (Chipewyans, Dakelh, etc.) and Southern Athabaskan. Importantly, the Central and West Siberian ancestry is almost exclusive to Na-Dene, and missing in other North or South American native ethnic groups, including Haida^20^, a group previously considered a divergent member of the Na-Dene language family^26^. Thus, the current consensus view of the Na-Dene language family^27^ and the distribution of recent Siberian ancestry match remarkably well. Although the small population sizes do not allow statistically valid comparisons, individuals with noticeable Saqqaq ancestry are likely more frequent among Northern Athabaskans as compared to Tlingit and Southern Athabaskans, the latter being mixed with southern Native Americans (Suppl. Fig. 1B).

We speculate that a migrating population, starting from Siberia around 6,500 YBP (the Syalakh culture), entering the New World around 4,800 YBP, and later mixing with First Americans, might have carried the Dene-Yeniseian languages^31–36^ into North America. This hypothetical language macrofamily unites multiple Na-Dene languages and Ket, the only surviving remnant of the Yeniseian family, once widespread in South and Central Siberia^33,37–40^. For a further description of the Dene-Yeniseian hypothesis and a review of lexicostatistical dating estimates see Suppl. Text 3. Although the Dene-Yeniseian macrofamily is not universally accepted among historical linguists^41,42^ (cf. Hamp^43^), and correlation of linguistic and genetic history is far from universal, the existence of the exclusive Siberian-Dene gene flow makes a genealogical relationship of the language families, either as the closest sister-groups^35^ or within a wider clade^42^, an attractive area of future research. Inferred age of the gene flow, 6,500-7,000 YBP, possibly corresponding to the split of Na-Dene and Yeniseian precursors in Siberia, is comparable to the age of the classic Indo-European language family^44,45^, suggesting that investigation of the Dene-Yeniseian connection lies within the reach of current methods in historical linguistics.

## Methods

### Sample collection and DNA extraction of 58 newly reported samples

Saliva samples of four Siberian ethnic groups (Enets, Kets, Nganasans, Selkups) were collected and DNA extractions were performed as described in Flegontov *et al.*^15^ Sampling locations and additional information is provided in Suppl. Table 3.

### Dataset preparation

In order to analyze rare allele sharing patterns, we composed a set of sequencing data covering Africa, Europe, Southeast Asia, Siberia, and the Americas: 1,206 individuals from 94 populations (Suppl. Table 1). Three sources were utilized to assemble the genome dataset: the Simons Genome Diversity Project^19^, Raghavan *et al*.^7^, and the 1000 Genomes Project^18^. We used variant calls generated in the respective publications, kept biallelic autosomal SNPs only and applied a filter based on a mappability mask^12^.

Additionally, we assembled two independent SNP datasets: see dataset compositions and filtration settings in Suppl. Tables 1 and 2. Initially, we obtained phased autosomal genotypes for large worldwide collections of Affymetrix HumanOrigins or Illumina SNP array data (Suppl. Table 2), using ShapeIt v.2.20 with default parameters and without a guidance haplotype panel^46^. Then we applied missing rate thresholds for individuals (<50% or <51%) and SNPs (<5%) using PLINK v.1.90b3.36^47^. For some analyses, unlinked SNPs were selected using linkage disequilibrium filtering with PLINK (Suppl. Table 2). Ten principal components (PC) were computed using PLINK on unlinked SNPs, and Euclidean distances defined as:

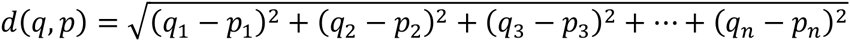

were calculated among individuals within populations (*q*_*n*_ and *p*_*n*_ refer to PCs from 1 to 10 in a population). We removed outliers according to the Euclidean distances, and populations having on average >5% of the Siberian ancestral component according to ADMIXTURE^48^ analysis (Suppl. Fig. 1), e.g. Finns and Russians, were excluded from the European and Southeast Asian meta-populations. In the case of the Illumina SNP array dataset, Na-Dene populations were exempt from PCA outlier removal and from removal of supposed relatives identified by Raghavan *et al.*^7^ It was done to preserve maximal diversity of Na-Dene and to ensure that both Dakelh individuals with sequencing data available would be included. Finally, we selected relevant meta-populations, generating datasets of 567-1,283 individuals further analyzed with ADMIXTURE^48^,

ChromoPainter v.1 and fineSTRUCTURE^17^, ChromoPainter v.2 and GLOBETROTTER^13^ (Suppl. Tables 1 and 2).

### Rare allele sharing statistics

We define the Allele Sharing Count between populations *A* and *B* (*ASC*_*AB*_ or *C*_*A,B*_) as the average number of sites at which an individual from population *A* shares a derived allele of frequency *k* with an individual from population *B:*

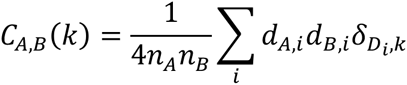

where *n* is the number of individuals in the populations, *d*_*A,i*_ stands for the number of derived alleles at site *i* in population *A,* and the term *δ*_*x,y*_ equals 0 if the total count of derived alleles in the dataset does not equal *k,* and is 1 otherwise. The sum across all sites *i* is normalized by the product of population sizes multiplied by four to give the average number of shared alleles between two randomly drawn haploid chromosome sets. Instead of counting derived alleles, in practice we counted non-reference alleles, which should not make a difference for low frequencies. To take care of variability in genome coverage across populations and of dataset-specific SNP calling biases, we calculated normalized allele sharing counts for populations *A* and *B,* dividing ASC*_AB_* by ASC_*AC*_, where population *C* is a distant outgroup. Because we assume that mutations occur as a Poisson process, the standard deviation of *ASC*_*AB*_ is defined as:

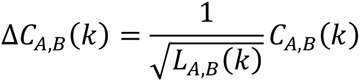

*L*_*A,B*_(*k*) is the number of sites *i*, at which derived alleles occur *k* times in the dataset. The standard deviation of *ASC*_*AB*_/*ASC*_*AC*_ is calculated using error propagation via partial derivatives:

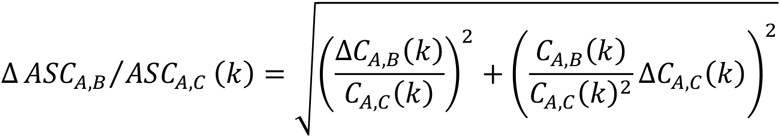

In practice, population *A* was an American population or ancient genome, population *B* was represented by Siberian or Arctic meta-populations, and population *C* – by Africans or Europeans (Suppl. Table 1). The resulting statistics are referred to as relative Siberian or Arctic allele sharing. Similar statistics were calculated for various Siberian and Arctic populations using the leave-one-out procedure. Allele sharing statistics were also calculated for private alleles and normalized by regular African or European ASCs. We called a shared allele private, or exclusively shared, if it was present in an American population and Siberians or members of the Arctic group, but missing in all other meta-populations (we did not condition on the presence of this allele in other Americans).

### Haplotype sharing statistics

Shared haplotype length (SHL_*AB*_) is defined as the total genetic length of DNA (in cM) that a given recipient individual *A*_*i*_ copies from a donor individual *B*_*j*_ under the model^13,17^. SHL_*AB*_ was computed in the all vs. all manner by ChromoPainter v.1^17^ running with default parameters. For each individual of a recipient population *A* (in practice an American individual), SHL_*AB*_ values were averaged across all individuals of a donor population *B* (the Siberian or Arctic meta-population), and then normalized by the haplotype sharing statistic SHL_*AC*_ for the European or African outgroup *C.* The resulting statistics SHL_*AB*_/SHL_*AC*_ are referred to as Siberian or Arctic relative haplotype sharing, and were visualized for separate individuals. Similar statistics were calculated for Siberian and Arctic individuals using the leave-one-out procedure at the population level.

Shared haplotype length (SHL_*AB*_) is defined as the total genetic length of DNA (in cM) that a given recipient individual *A*_*i*_ copies from a donor individual *B*_*j*_ under the model^13,17^. SHL_*AB*_ was computed in the all vs. all manner by ChromoPainter v.1^17^ running with default parameters. For each individual of a recipient population *A* (in practice an American individual), SHL_*AB*_ values were averaged across all individuals of a donor population *B* (the Siberian or Arctic meta-population), and then normalized by the haplotype sharing statistic SHL_*AC*_ for the European or African outgroup *C.* The resulting statistics SHL_*AB*_/SHL_*AC*_ are referred to as Siberian or Arctic relative haplotype sharing, and were visualized for separate individuals. Similar statistics were calculated for Siberian and Arctic individuals using the leave-one-out procedure at the population level.

### Dating admixture events using haplotype sharing statistics

We used GLOBETROTTER^13^ to infer and date up to two admixture events in the history of Na-Dene populations. To detect subtle signals of admixture between closely related partners, we followed the ‘regional’ analysis protocol of Hellenthal *et al.*^13^ Using ChromoPainter v.2^13^, Na-Dene chromosomes were ‘painted’ as a mosaic of haplotypes derived from donor populations or meta-populations: the Saqqaq ancient genome, Siberian Arctic populations, American Arctic populations, northern North Americans, other Americans, core Siberians, Siberians with ANE ancestry, Southeast Asians, Europeans. Na-Dene individuals were considered as haplotype recipients only, while other populations or meta-populations were considered as both donors and recipients. That is different from the ChromoPainter v.1 approach, where all individuals were considered as donors and recipients of haplotypes at the same time, and only self-copying was forbidden.

Painting samples for Na-Dene (the target population) and ‘copy vectors’ for other (meta)populations called ‘surrogates’ served as an input of GLOBETROTTER, which was run according to section 6 of the instruction manual of May 27, 2016. The following settings were used: no standardizing by a “NULL” individual (null.ind 0); five iterations of admixture date and proportion/source estimation (num.mixing.iterations 5); at each iteration, any surrogates that contributed ≤ 0.1% to the mixture describing the target population were removed (props.cutoff 0.001); the x-axis of coancestry curves spanned the range from 0 to 50 cM (curve.range 1 50), with bins of 0.1 cM (bin.width 0.1). Confidence intervals (95%) for admixture dates were calculated based on 100 bootstrap replicates with options null.ind 0 and num.admixdates.bootstrap 2 (fitting two dates when performing bootstrapping). Generation time of 29 years was used in all dating calculations^7^.

The GLOBETROTTER software is able to date no more than two admixture events^13^, therefore we had to reduce the complexity of original Na-Dene populations that likely experienced more than two major waves of admixture. For that purpose, only a subset of Na-Dene individuals was used for the GLOBETROTTER analysis: individuals demonstrating a signal of Paleo-Eskimo admixture and a low level of European ancestry according to Siberian and Arctic haplotype sharing statistics with the European normalizer. In practice, Na-Dene individuals lying in the area of the two-dimensional plot occupied by simulated mixtures of the 1^st^ and 2^nd^, but not the 1^st^ and 3^rd^ migration waves (Fig. 4), were treated as one target population. This definition of a ‘target’ population was used with meta-populations as haplotype donors. To increase the amount of data when separate populations were used for calculating coancestry curves, we included 8 additional Chipewyan individuals with evidence of low-level Paleo- or Neo-Eskimo admixture, i.e. lying in the area of the two-dimensional plot where the two clusters of simulated mixtures overlap (Fig. 4).

### Rarecoal analysis

We used the Rarecoal program (https://github.com/stschiff/rarecoal) to fit demographic models to meta-populations, iteratively adding one population at a time (Suppl. Text 1). We started with a tree connecting Europeans, Southeast Asians, and Native Americans into a simple tree without admixture, and used “rarecoal mcmc” to infer maximum likelihood branch population sizes and split times. We then iteratively added additional populations, and after each addition, we re-optimized the tree and inspected the fits of the model to the data. When we saw a significant deviation between model and data, we added admixture edges, informed by the under- or over-estimation of a particular sharing pattern (Suppl. Text 1).

After Rarecoal’s inference, we rescaled time and population size parameters to years and real effective population size using a mutation rate of 1.25'10^-8^ per site per generation, and a generation time of 29 years^7^. The final model, as shown in Figure 5, was then also simulated using the SCRM simulator^49^, and we verified that Rarecoal was able to infer the true parameters after simulation.

In order to map the two ancient genomes, Saqqaq and Clovis, onto the tree, we restricted the analysis to variants between allele counts 2 and 4. We excluded singletons, because they are highly enriched for false positives in ancient genomes, and mainly used for population size estimation, which we are less interested in in the case of ancient samples^12^. We used “rarecoal find” to evaluate the likelihood for merging onto the tree at all branches and all times (after the date of the sample).

### ADMIXTURE analysis

The ADMIXTURE software^48^ implements a model-based Bayesian approach that uses block-relaxation algorithm in order to compute a matrix of ancestral population fractions in each individual (*Q*) and infer allele frequencies for each ancestral population (*P*). A given dataset is usually modeled using various numbers of ancestral populations *(K).* We ran ADMIXTURE on HumanOrigins-based and Illumina-based datasets of unlinked SNPs (Suppl. Table 2) using 10 to 25 and 5 to 20 *K* values, respectively. One hundred analysis iterations were generated with different random seeds. The best run was chosen according to the highest likelihood. An optimal value of *K* was selected using 10-fold cross-validation (CV).

### fineSTRUCTURE: PCA and clustering

We used fineSTRUCTURE v.2.0.7 with default parameters to analyze the output of ChromoPainter v.1^17^. Clustering trees of individuals were generated by

fineSTRUCTURE based on counts of shared haplotypes^17^. The clustering trees and coancestry matrices were visualized using fineSTRUCTURE GUI v.0.1.0^17^. Finally, PCA was generated based on counts of shared haplotypes and visualized using R.

## Acknowledgements

We are grateful to all researchers that shared their data: David Reich, Nick Patterson, Iain Mathieson, Swapan Mallick, Maanasa Raghavan, Simon Rasmussen, and Eske Willerslev. We also thank David Reich for helpful comments and for curating the newly reported HumanOrigins genotyping data. P.F. was supported by the Institution Development Program of the University of Ostrava and by EU structural funding Operational Programme Research and Development for Innovation, project No. CZ.1.05/2.1.00/19.0388.

## Author contributions

P.F. and S.S. have designed the study, analyzed the data and written the manuscript; N.E.A. and P.C. have analyzed the data and prepared the figures and tables; E.J.V. has contributed the sections dealing with Na-Dene linguistics and archaeology; and J.K. was involved in sample genotyping and in manuscript preparation.

